# Transcriptome Deconvolution of Heterogeneous Tumor Samples with Immune Infiltration

**DOI:** 10.1101/146795

**Authors:** Zeya Wang, Shaolong Cao, Jeffrey S. Morris, Jaeil Ahn, Rongjie Liu, Svitlana Tyekucheva, Fan Gao, Bo Li, Wei Lu, Ximing Tang, Ignacio I. Wistuba, Michaela Bowden, Lorelei Mucci, Massimo Loda, Giovanni Parmigiani, Chris C. Holmes, Wenyi Wang

**Affiliations:** Department of Bioinformatics and Computational Biology, The University of Texas MD Anderson Cancer Center, Houston, TX, United States; Department of Statistics, Rice University, Houston, TX, United States; Department of Biostatistics, The University of Texas MD Anderson Cancer Center, United States; Department of Biostatistics and Bioinformatics, Georgetown University, Washington, DC, United States; Department of Biostatistics and Computational Biology, Dana Farber Cancer Institute, Boston, MA, United States; Department of Statistics, Harvard University, Cambridge, MA, United States; Department of Translational Molecular Pathology, The University of Texas MD Anderson Cancer Center, Houston, TX, United States; Department of Oncologic Pathology, Dana Farber Cancer Institute, Boston, MA, United States; Department of Epidemiology, Harvard T.H. Chan School of Public Health, Boston, MA, United States; Department of Pathology, Brigham and Women’s Hospital and Harvard Medical School, Boston, MA, United States; Department of Biostatistics, Harvard T.H. Chan School of Public Health, Boston, MA, United States; Department of Statistics, University of Oxford, Oxford, United Kingdom

**Keywords:** tumor heterogeneity, RNA-seq data, tumor-stroma-immune interaction, head and neck squamous cell carcinoma, statistical models, computational tool

## Abstract

Transcriptomic deconvolution in cancer and other heterogeneous tissues remains challenging. Available methods lack the ability to estimate both component-specific proportions and expression profiles for individual samples. We present *DeMixT*, a new tool to deconvolve high dimensional data from mixtures of more than two components. *DeMixT* implements an iterated conditional mode algorithm and a novel gene-set-based component merging approach to improve accuracy. In a series of experimental validation studies and application to TCGA data, *DeMixT* showed high accuracy. Improved deconvolution is an important step towards linking tumor transcriptomic data with clinical outcomes. An R package, scripts and data are available: https://github.com/wwylab/DeMixT/.

## Background

Heterogeneity of malignant tumor cells adds confounding complexity to cancer treatment. The evaluation of individual components of tumorsamples is complicated by the tumor-stroma-immune interaction. Anatomical studies of the tumor-immune cell contexture have demonstrated that it primarily consists of a tumor core, lymphocytes and the tumor microenvironment [8,19]. Further research supports the association of infiltrating immune cells with clinical outcomes for individuals with ovarian cancer, colorectal cancer and follicular lymphoma [6,9,26]. The use of experimental approaches such as laser-capture microdissection (LCM) and cell sorting is limited by the associated expense and time. Therefore, understanding the heterogeneity of tumor tissue motivates a computational approach to integrate the estimation of type-specific expression profiles for tumor cells, immune cells and the tumor microenvironment. Most commonly available deconvolution methods assume that malignant tumor tissue consists of two distinct components, epithelium-derived tumor cells and surrounding stromal cells [2,11]. Other deconvolution methods for more than two compartments require knowledge of cell-type-specific gene lists [14], i.e., reference genes, with some of these methods applied to estimate subtype proportions within immune cells [12,18]). Therefore, there is still a need for methods that can jointly estimate the proportions and compartment-specific gene expression for more than two compartments in each tumor sample.

The existing method ISOpure [20] may address this important problem. However, ISOpure assumes a linear mixture of raw expression data and represents noncancerous profiles in the mixed tissue samples by a convex combination of all the available profiles from reference samples. A drawback of this modeling approach is that the variance for noncancerous profiles is not compartment-specific, therefore: 1) the variances that are needed for estimating sample- and compartment-specific expressions cannot be estimated; and 2) not accounting for sample variances can result in large bias in the estimated mixing proportions and mean expressions. As we aim to address the need for having both the gene-specific variance parameters and two unknown mixing proportions per sample in the 3-component scenario, our previous heuristic search algorithm developed for 2 components [2] is inadequate for the computation.

We have developed a new computational tool, *DeMixT*, to accurately and efficiently estimate the desired high-dimensional parameters, in a linear additive model that accounts for variance in gene expression levels in each compartment (**Figure 1a**). The corresponding R package for *DeMixT* is freely available for downloading at https://github.com/wwylab/DeMixr.

**Figure 1.**
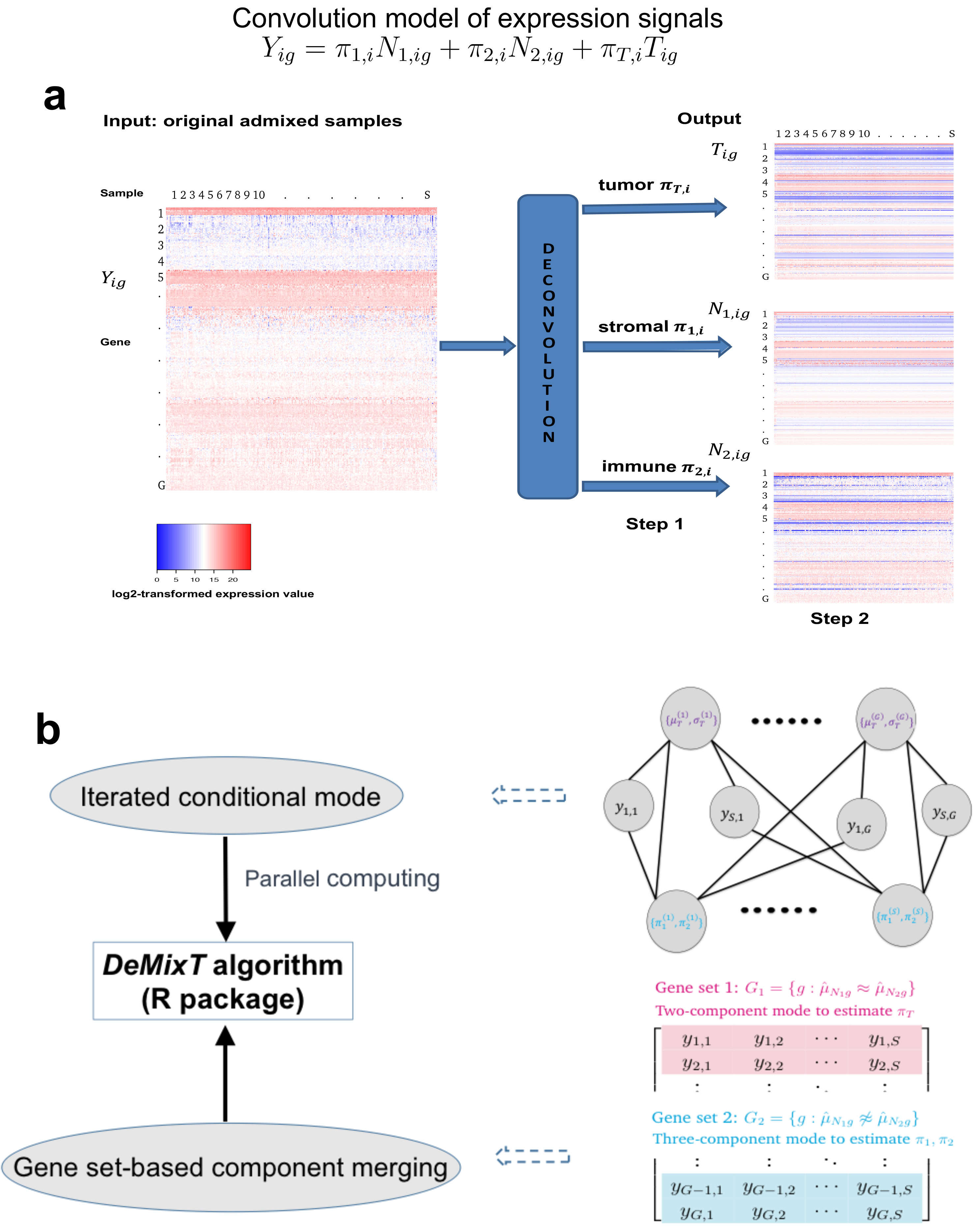
The model and algorithm of *DeMixT*. (a) *DeMixT* performs a three-component deconvolution to output tissue-specific proportions and isolated expression matrices of tumor (T-component), stromal (*N*_1_-component) and immune cells (*N*_2_-component). Heatmaps of expression levels correspond to the original admixed samples, the deconvolved tumor component, stromal component and immune component. (b) The *DeMixT*-based parameter estimation is achieved by using the iterated conditional modes (ICM) algorithm and a gene-set-based component merging (GSCM) approach. The top graph describes the conditional dependencies between the unknown parameters, which can be assigned to two groups: genome-wise parameters (on the top row, with red superscript), and sample-wise parameters (on the bottom row, with blue superscript). They are connected by edges, which suggest conditional dependencies. The nodes on the top row that are not connected are independent of each other when conditional on those on the bottom row, and vice versa. Because of the conditional independencies, we implemented parallel computing to substantially increase the computational efficiency. The bottom graph illustrates the GSCM approach, which first runs a two-component deconvolution on gene set *G*_1_ (red), where 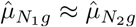 in order to estimate *π*_*T*_, and then runs a three-component deconvolution on gene set *G*_2_ (blue), where 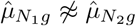 and *π*_*T*_ is given by the prior step, in order to estimate *π*_1_ and *π*_2_.

## Results

*The DeMixT model and algorithm.* Here, we summarize our convolution model as follows (**Figure 1a**; see further details in Methods). The observed signal *Y*_*ig*_ is written as *Y*_*ig*_ = **π**_1,*i*_*N*_1,*ig*_ + *π*_2,*i*_*N*_2,*ig*_ + (1 − *π*_1,*i*_ − *π*_2,*i*_)*T*_*ig*_ for each gene *g* and each sample *i*, where *Y*_*ig*_ is the expression for the observed mixed tumor samples, and *N*_1,*ig*_, *N*_2,*ig*_ and *T*_*ig*_ represent unobserved raw expression values from its constituents. We assume that *N*_1,*ig*_, *N*_2,*i*_ and *T*_*ig*_, each follow a log_2_-normal distribution with compartment-specific means and variances [2,15]. The *N*_1_-component and the *N*_2_-component are the first two components, the distributions of which need to be estimated from available reference samples, and *π*_1,*i*_ and *π*_2,*i*_ are the corresponding proportions for sample *i*. The last component is the T-component, the distribution of which is unknown. In practice, the T-component can be any of the following three cell types: tumor, stromal or immune cells. For inference, we calculate the full likelihood and search for parameter values that maximize the likelihood. Our previously developed heuristic search algorithm [2] for a two-component model is inadequate for a three-component model which is exponentially more complex: 1) There are two degrees of freedom in the mixing proportions, which is unidentifiable in a large set of genes that are not differentially expressed between any two components; and 2) In each iteration in the parameter search, we need to perform tedious numerical double integrations in order to calculate the full likelihood. The *DeMixT* algorithm introduced two new elements that helped ensure the estimation accuracy and efficiency (**Figure 1b**). We first apply an optimization approach, iterated conditional modes (ICM) [4], which cyclically maximizes the probability of each set of variables conditional on the rest, for which we have observed rapid convergence [4] to a local maximum. The ICM framework further enables parallel computing, which helps compensate for the expensive computing time used in the repeated numerical double integrations. However this is not sufficient for accurate parameter estimation. We observed that including genes that are not differentially expressed between the *N*_1_ and *N*_2_ components in the 3-component deconvolution can introduce large biases in the estimated *π*_1_ and *π*_2_ (**Supplementary Figure 1**), while the *π*_*T*_ estimation is little unaffected. We therefore introduce a novel gene-set-based component merging approach (GSCM) (**Figure 1b**). Here, we first select gene set 1, where *μ*_1,*g*_ ≈ *μ*_2,g_, and run the 2-component model to estimate *π*_*T,i*_. Then we select gene set 2, where 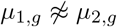, and run the 3-component model with fixed *π*_*T*_ from the equation above, to estimate {*π*_1,*i*_, *π*_2,*i*_}. Our goal is to avoid searching in the relatively flat regions of the full likelihood (model unidentifiable, **Supplementary Figure 2**) and focus on regions where the likelihood tends to be convex. Using this approach, we not only improve the estimation accuracy, but also further reduce the computing time as only a small part of the entire parameter space needs to be searched.

*Validation using data with known truth.* We validated *DeMixT* in two datasets with known truth in proportions and mean expressions: a publicly available microarray dataset [21] generated using mixed RNA from rat brain, liver and lung tissues in varying proportions; and an RNA-seq dataset generated using mixed RNA from three cell lines, lung adenocarcinoma (H1092), cancer-associated fibroblasts (CAFs) and tumor infiltrating lymphocytes (TILs).

We used GSE19830 [21] as our first dataset for benchmarking. This microarray experiment was designed for expression profiling of samples from *Rattus norvegicus* with the Affymetrix Rat Genome 230 2.0 Array, including 30 mixed samples of liver, brain and lung tissues in 10 different mixing proportions with three replicates (Supplementary Table 1). To run *DeMixT*, we used the samples with 100% purity to generate the reference profiles for the *N*_1_-component, *N*_2_-component and *T*-component, respectively. We ran the deconvolution for the 30 mixed samples as three scenarios, respectively assuming the liver, brain and lung tissues to be the unknown *T*-component tissue. To generate the second dataset in RNA-seq, we performed a mixing experiment, in which we mixed mRNAs from the three cell lines: lung adenocarcinoma in humans (H1092), CAFs and tumor infiltrating lymphocytes (TILs), at different proportions to generate 32 samples, including 9 samples that correspond to three repeats of a pure cell line sample for the three cell lines (Supplementary Table 2). The RNA amount of each tissue in the mixture samples was calculated on the basis of real RNA concentrations tested in the biologist’s lab. We assessed our deconvolution approach through a number of statistics, e.g., concordance correlation coefficients (CCCs), root mean squared errors (RMSEs), and a summary statistics for measuring the reproducibility of the estimated *π* across scenarios when a different component is unknown (see Methods). We showed that *DeMixT* performed well and outperformed ISOpure in terms of accuracy and reproducibility (**Figures 2a-b**; see **Supplementary Notes** for further details, **Supplementary Figures 3-6**, **Table 1**, **Supplementary Tables 3-6**).

**Figure 2.**
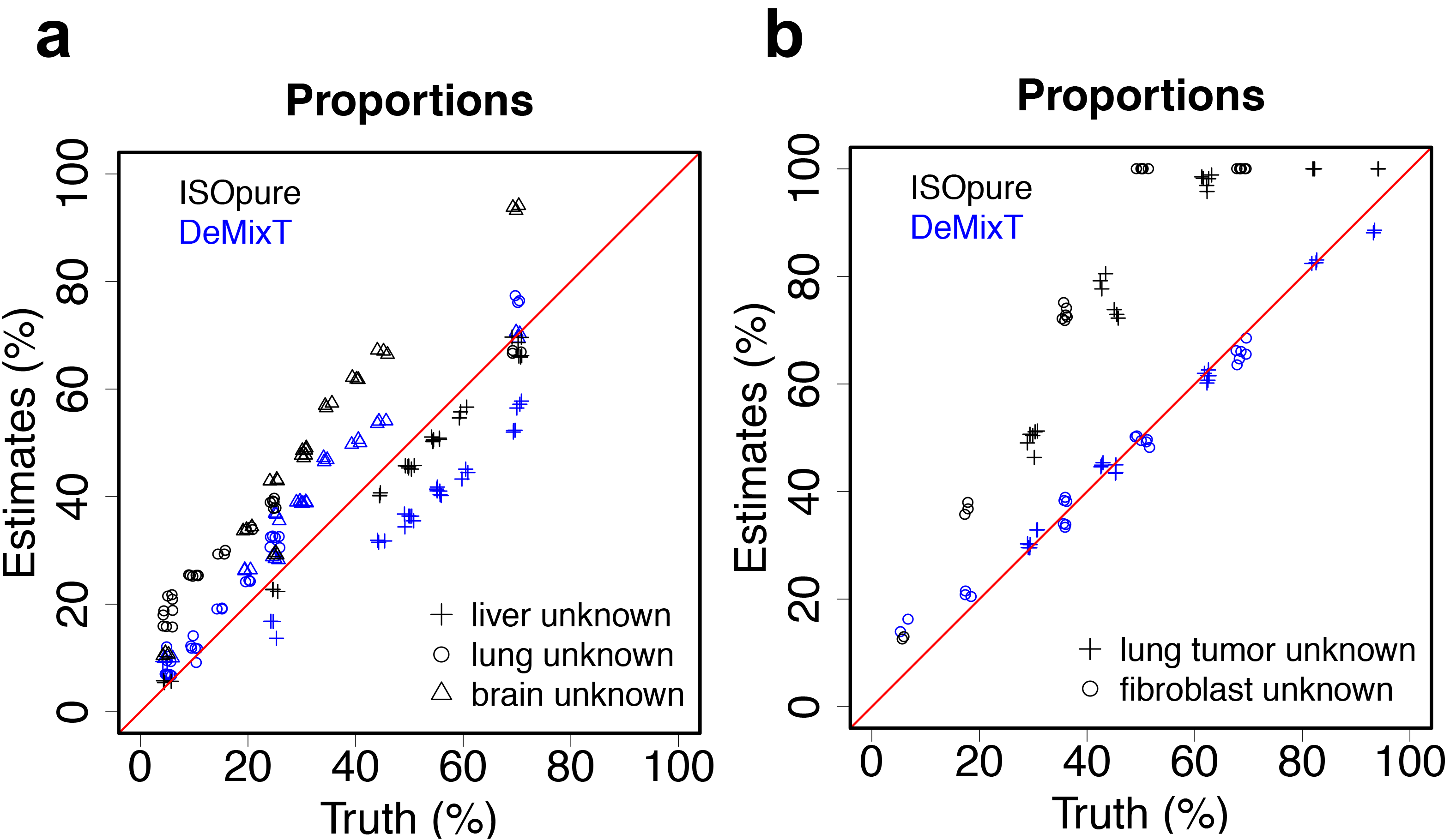
Validation results using microarray and RNA-seq data from tissue and cell line mixture experiments. (a) A scatter plot of estimated tissue proportions vs. the truth when the liver (cross), brain (triangle), or lung (rectangle) tissue is assumed to be the unknown tissue in the microarray experiments mixing the three; estimates from ISOpure are also presented. (b) Scatter plot of estimated tissue proportions vs. the truth when either lung tumor (cross) or fibroblast (rectangle) cell lines are assumed to be the unknown tissue in the RNA-seq experiments mixing lung tumor, fibroblast and lymphocyte cell lines.

**Table 1.** Measures of reproducibility for estimated proportions across different scenarios in the GSE19830 dataset and the mixed cell line RNA-seq dataset.

| Estimated Tissue | *DeMixT* | ISOpure |
| --- | --- | --- |
| Brain | 0.03 | 0.10 |
| Lung | 0.03 | 0.08 |
| Liver | 0.03 | 0.07 |
| H1092 | 0.05 | 0.40 |
| CAF | 0.06 | 0.41 |
| TIL | 0.02 | 0.02 |
| H1092, lung tumor adenocarcinoma; CAF, cancer-associated fibroblast; TIL, tumor infiltrating lymphocytes |  |  |

*Validation using LCM data*. We then applied *DeMixT* to a “gold standard” validation dataset from real tumor tissue that has known proportions, mean expressions and individual component-specific expressions. This dataset (GSE97284) was generated at Dana Farber Cancer Institute through LCM experiments on tumor samples from patients with prostate cancer. It consists of 25 samples of isolated tumor tissues, 25 samples of isolated stromal tissues and 23 admixture samples [24]. LCM was performed on formalin-fixed paraffin embedded (FFPE) tissue samples from 23 prostate cancer patients, and microarray gene expression data were generated using the derived and the matching dissected stromal and tumor tissues (GSE97284 [23]). Due to the low quality of the FFPE samples, we selected a subset of probes (see Methods), and ran *DeMixT* under a two-component mode. *DeMixT* obtained concordant estimates of the tumor proportions when the proportion of the stromal component was unknown and when the proportion of tumor tissue was unknown (CCC=0.87) (**Figure 3a**). *DeMixT* also tended to provide accurate component-specific mean expression levels (**Figures 3b-c** and **Supplementary Figure 7**) and yielded standard deviation estimates that are close to those from the dissected tumor samples (**Supplementary Figure 8**). As a result, the *DeMixT* individually deconvolved expressions achieved high CCCs (mean= 0.96) for the tumor component (**Figure 3d** and **Supplementary Figure 9**). The expressions for the stromal component were more variable than those for a common gene expression dataset, hence both *DeMixT* and ISOpure gave slightly biased estimates of the means and standard deviations.

**Figure 3.**
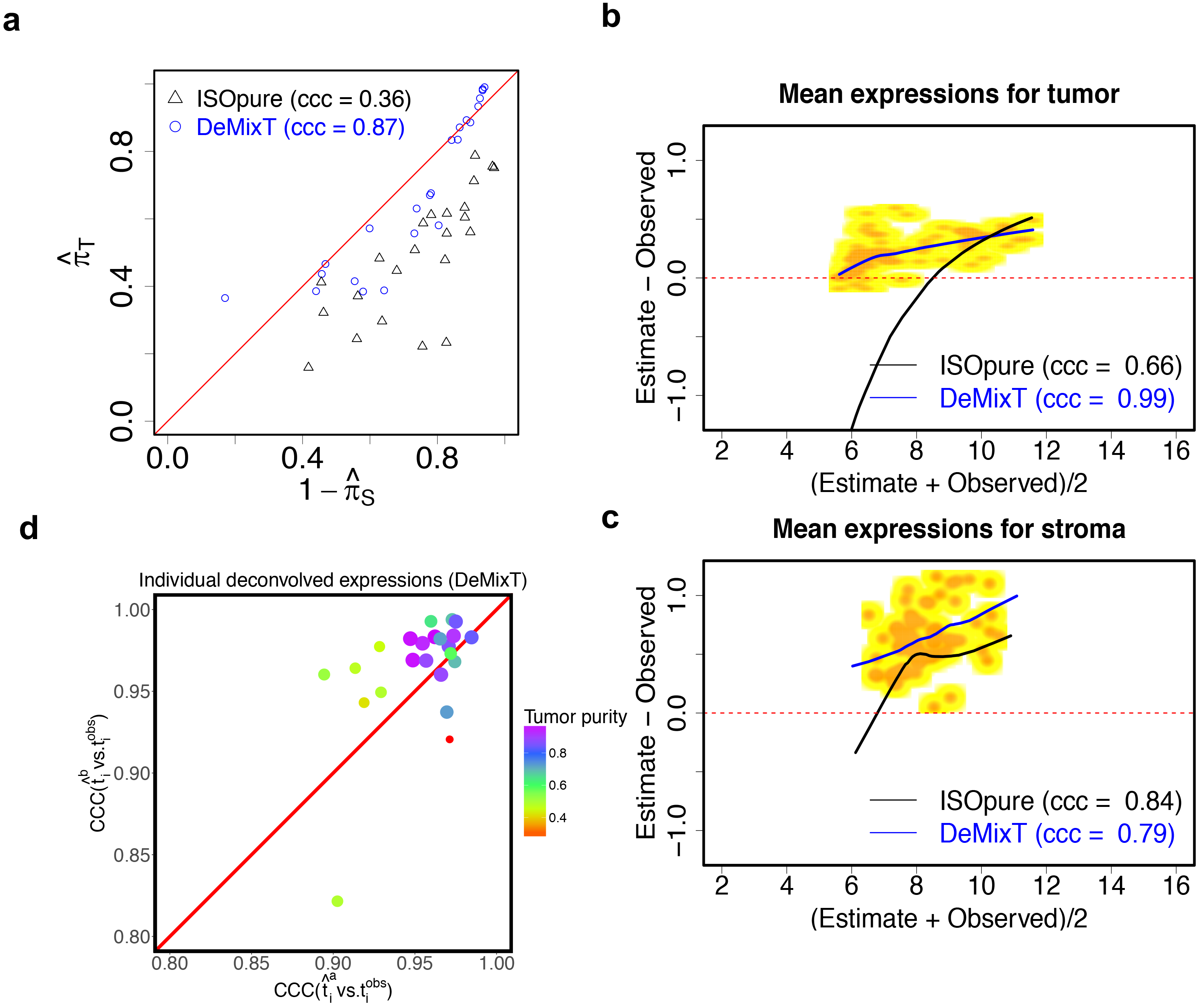
Analyses of real data using *DeMixT* through validation using LCM data in prostate cancer. (a) Scatter plot of estimated tumor proportions versus 1-estimated stromal proportions; estimates from *DeMixT* (blue) are compared with those from ISOpure (black). (b)-(c) Smoothed scatter MA plots between observed and deconvolved mean expression values from *DeMixT* for the tumor and stromal components, respectively (yellow for low values and orange for high values). The lowess smoothed curves for *DeMixT* are shown in black and those for ISOpure in blue. (d) Scatter plot of concordance correlation coefficient (CCC) between individual deconvolved expression profiles 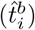 and observed values 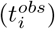 for 23 LCM matching prostate cancer samples. Superscript *a*: stromal component is represented by reference samples; *b*: tumor component is represented by reference samples. Color gradient of each point corresponds to the estimated tumor proportion.

*Application to The Cancer Genome Atlas (TCGA) head and neck squamous cell carcinoma (HNSCC) data*. A recent study of HNSCC showed that the infiltration of immune cells, both lymphocytes and myelocytes, is positively associated with viral infection in virus-associated tumors [13]. We downloaded HNSCC RNA-seq data from TCGA data portal [17] and ran *DeMixT* for deconvolution. We normalized the expression data with the total count method and filtered out genes with zero count in any sample. There was a total of 44 normal tissue and 269 tumors samples in the HNSCC dataset. We collected the infection information of human papillomavirus (HPV) infection status for the HNSCC samples. Samples were classified as HPV-positive (HPV+) using an empiric definition of the detection of > 1000 mapped RNA-Seq reads, primarily aligning to viral genes E6 and E7, which resulted in 36 HPV+ samples [17]. Since only reference samples for the stromal component are available from TCGA (i.e., 44 normal samples and 269 tumor samples), we devised an analytic pipeline for *DeMixT* to run successfully on the HNSCC samples (for details, see Methods and **Supplementary Figure 10**). In brief, we first used data from the HPV+ tumors to derive reference samples for the immune component, and then ran the three-component *DeMixT* on the entire dataset to estimate the proportions for both HPV-negative (HPV−) and HPV+ samples. For all tumor samples, we obtained the immune (mean = 0.22, standard deviation = 0.10), the tumor (mean = 0.64, standard deviation = 0. 13), and the stromal proportions (mean = 0.14, standard deviation = 0.07; see **Figure 4a**). The distribution of stromal proportions seems independent, whereas the tumor and immune proportions are inversely correlated. As expected, HPV+ tumor samples had significantly higher immune proportions than those that tested as HPV−, [7,13] (p-value = 2e-8; **Figures 4a-b** and **Supplementary Figure 11**). To further evaluate the performance of our deconvolved expression levels, we performed differential expression tests for immune versus stromal tissue and immune versus tumor tissue, respectively, on 63 infiltrating immune cell-related genes (CD and HLA genes). For example, **Figure 4c** illustrates that the deconvolved expressions were much higher in the immune component than in the other two components for three important immune marker genes, CD4, CD14, and HLA-DOB. Overall, 51 out of 63 genes were significantly more highly expressed in the immune component than in the other two components (adjusted p-values are listed in **Supplementary Data 1**; also see **Figure 4d**).

**Figure 4.**
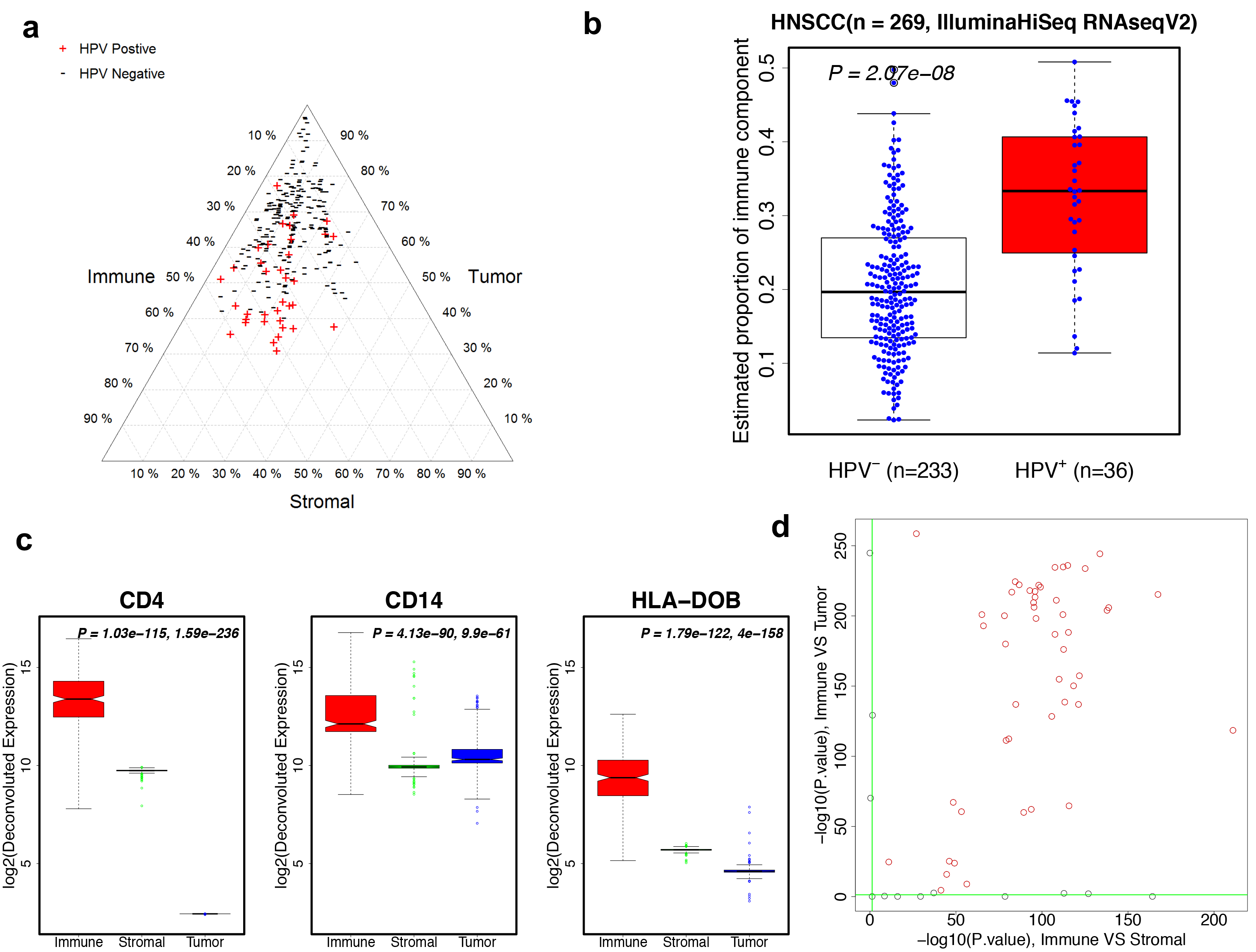
Analyses of real data using *DeMixT* through application to TCGA RNA-seq data in HNSCC. (a) Triangle plot of estimated proportions (%) of the immune component (left), tumor component (right) and stromal component (bottom) in HNSCC data; “+” and “−” corresponds to the HPV status. (b) Boxplots of estimated immune proportions for HNSCC samples in the test set display differences between HPV+ (red) and HPV− (white) samples. (c) Boxplots of log2-transformed deconvolved expression profiles for three important immune genes (CD4, CD14, HLA-DOB) in the test set of HNSCC samples. Red: immune component; green: stromal component; blue: tumor component. P-values of differential tests are at top right corner for each gene: the first p-value is for immune vs. stromal component; second p-value is for immune vs. tumor component. (d) Scatter plot of negative log-transformed p-values for comparing deconvolved expression profiles between immune component and the other two components of 63 immune cell-related genes. The x-axis: immune component vs. stromal component; y-axis: immune component vs. tumor component. Genes in red are significant in both comparisons. Green horizontal and vertical lines: cutoff value for statistical significance.

## Conclusions

In this work, we have presented a novel statistical method and software, *DeMixT* (R package at https://github.com/wwylab/DeMixT), for dissecting a mixture of tumor, stromal and immune cells bsed on the gene expression levels, and providing an accurate solution. Our method allows us to simultaneously estimate both cell-type-specific proportions and reconstitute patient-specific gene expression levels with little prior information. Distinct from the input data of most other deconvolution methods, our input data consist of gene expressions from 1) observed mixtures of tumor samples and 2) a set of reference samples from *p* − 1 compartments (where *p* is the total number of compartments). Our output data provides the mixing proportions, the means and variances of expression levels for genes in each compartment, as well as the expression levels for each gene in each compartment and each sample. The full gene-compartment sample-specific output allows for the application of all pipelines previously developed for downstream analyses, such as clustering and feature selection methods in cancer biomarker studies, still applicable to the deconvolved gene expressions. We achieved this output by modeling compartment-specific variance and addressing the associated inferential challenges. *DeMixT* addresses the transcriptomic deconvolution in two steps. In the first step, we estimate the mixing proportions and the gene-specific distribution parameters for each compartment, using the ICM method, which can quickly converge. We further proposed a novel gene-set-based component merging (GSCM) approach and integrated it with ICM for the three-component deconvolution, in order to substantially improve model identifiability and computational efficiency. In the second step, we reconstitute the expression profiles for each sample and each gene in each compartment based on the parameter estimates from the first step. The success of the second step relies largely on the success of the first. We have overcome the otherwise significant computational burden for searching the high-dimensional parameter space and numerical double integration, due to our explicit modeling of variance, through parallele computing and gene set-based componment merging. On a PC with a 3.07 GHz Intel Xeon processor with 20 parallel threads, *DeMixT* takes 14 minutes to complete the full three-component deconvolution task of a dataset consisting of 50 samples and 500 genes (see **Table 2**). Our method can be applied to other data types such as proteomic data. It can be extended to *p* > 3.

**Table 2.** Computing time for *DeMixT*. *DeMixT* was run on a simulated dataset consisting of 50 samples and 500 genes using 2 or 20 threads. Of all genes, 400 belong to gene set 1 (G_1_) and the remaining 100 belong to gene set 2 (G_2_), as defined in our gene-set-based component merging approach (see Figure 1b).

|  | w/o CM | w/ CM |  |  |
| --- | --- | --- | --- | --- |
| | Total | Two-component step: $G_1$ | Three-component step: $G_2$ | Total |
| 2 threads | 16.1 h | 37 min | 48 min | 85min |
| 20 threads | 2.5 h | 6 min | 8 min | 14min |

We have used a series of experimental datasets to validate the performance of *DeMixT*. These datasets were generated from a mixture of normal tissues, a mixture of human cell lines, and LCM of FFPE tumor samples. *DeMixT* succeeded in recapitulating the truth in all datasets. We further demonstrated the tumor-stroma-immune deconvolution by *DeMixT* using TCGA HNSCC data. We were able to correlate our immune proportion estimates with the available HPV infection status in HNSCC, as is consistent with previous observations that a high level of immune infiltration appears with viral infection in cancer [12]. For this dataset, *DeMixT* is the first to provide a triangular view of tumor-stroma-immune proportions (**Figure 4a**), the interesting dynamics of which may shed new light on the prognosis of HNSCC.

Gene selection is important for the success of deconvolution in both 2-component and 3-component models for *DeMixT*, and for any other methods. In a 2-component setting, we observed that both variances and mean differences in expression levels between the two components for each gene can affect how accurately the mixing proportions are estimated, while not all genes are needed for the proportion estimation. We therefore suggested selecting genes that have moderate variances and large differences between two components to estimate proportions. In a 3-component setting, using the gene-set-based component merging (GSCM) approach to reduce to a pseudo-2-component problem, we are able to apply a similar strategy. Currently our gene selection and GSCM strategy follow the principle of focusing on a subspace of the high-dimensional parameters for model identifiability, but are ad hoc and may need modification across datasets. Our future work will be to systematically evaluate the impact of each set of high-dimensional parameters on the full likelihood underlying our convolution model, and search for a unified gene selection method for the deconvolution of datasets that range over a wide spectrum of biological phenomena.

The reference gene-based deconvolution is popular for estimating immune subtypes within immune cells [14, 18]. Our method does not require reference genes which we consider as difficult to obtain for the tumor component; however *DeMixT* can take reference genes when available. With the reference sample approach, we assume the first *p*-1 compartments in the observed mixture to be similar to those in reference samples, while the remaining compartment is unknown, and so it may end up capturing most of the heterogeneity. The reference samples can be derived from historical patient data or from the corresponding healthy tissues, such as data from GTEx [16] (e.g. RNA-seq data from sun-exposed skin as reference samples for melanoma, unpublished results). Furthermore, each of the three components may contain more than one type of cell, in particular, the immune component. It was reported that although heterogeneous, the relative proportions of immune cell subtypes within the immune component are consistent across patient samples [10], which supports our approach that models the pooled immune cell population using one distribution. Further development of *DeMixT* will account for subpopulations of immune cells in the convolution model, which will be a natural extension of the current model. Finally, the performance of *DeMixT* will be optimized when the data analysis practice is linked with the cancer-specific biological knowledge.

## Methods

### Model

Let *Y*_*ig*_ be the observed expression levels of the raw measured data from clinically derived malignant tumor samples for gene *g*, *g* = 1, ···, *G* and sample *i*, *i* = 1, ···, *S*. *G* denotes the total number of probes/genes and *S* denotes the number of samples. The observed expression levels for solid tumors can be modeled as a linear combination of raw expression levels from three components:

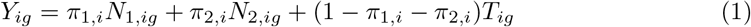

Here *N*_1,*ig*_, *N*_2,*ig*_ and *T*_*ig*_ are the unobserved raw expression levels from each of the three components. We call the two components for which we require reference samples the *N*_1_-component and the *N*_2_-component. We call the unknown component the T-component. We let *π*_1,*i*_ denote the proportion of the *N*_1_-component, *π*_2,*i*_ denote the proportion of the *N*_2_-component, and 1 − *π*_1,*i*_ − *π*_2,*i*_ denote the proportion of the T-component. We assume that the mixing proportions of one specific sample remain the same across all genes.

Our model allows for one component to be unknown, and therefore does not require reference profiles from all components. A set of samples for *N*_1,*ig*_ and *N*_2,*ig*_, respectively, needs to be provided as input data. This three-component deconvolution model is applicable to the linear combination of any three components in any type of material. It can also be simplified to a two-component model, assuming there is just one *N*-component. For application in this paper, we consider tumor (*T*), stromal (*N*_1_) and immune components (*N*_2_) in an admixed sample (*Y*).

Following the convention that log_2_-transformed microarray gene expression data follow a normal distribution, we assume that the raw measures 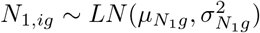, 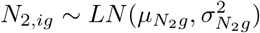 and 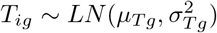, where LN denotes a log_2_-normal distribution and 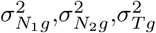 reflect the variations under log_2_-transformed data [2,15]. Consequently, our model can be expressed as the convolution of the density function for three log_2_-normal distributions. Because there is no closed form of this convolution, we use numerical integration to evaluate the complete likelihood function (see the full likelihood in the **Supplementary Materials**).

### The *DeMixT* algorithm for deconvolution

*DeMixT* estimates all distribution parameters and cellular proportions and reconstitutes the expression profiles for all three components for each gene and each sample, as shown in equation (1). The estimation procedure (summarized in **Figure 1b**) has two main steps as follows.

1. Obtain a set of parameters 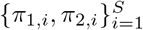, 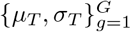 to maximize the complete likelihood function, for which 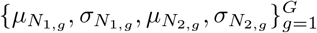 were already estimated from the available unmatched samples of the *N*_1_ and *N*_2_ component tissues. This step is described in further details below in parameter estimation and the GSCM approach.
2. Reconstitute the expression profiles by searching each set of {*n*_1,*ig*_, *n*_2,*ig*_} that maximizes the joint density of *N*_1,*ig*_, *N*_2,*ig*_ and *T*_*ig*_

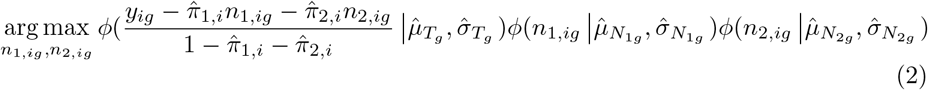

where *ϕ*(.|**μ*,*σ*^2^*) is a log2-normal distribution density with location parameter *μ* and scale parameter *σ*.

In step 2, we combined the golden section search method with successive parabolic interpolations to find the maximum of the joint density function with respect to *n*_1,*ig*_ and *n*_2,*ig*_ that are positively bounded and constrained by 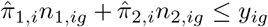. The value of *t*_*ig*_ is solved as 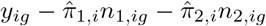.

### Parameter estimation using iterated conditional modes (ICM)

In step 1, the unknown parameters to be estimated can be divided into two groups: gene-wise parameters, 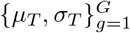, and sample-wise parameters, 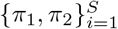. These two groups of parameters are conditionally independent (**Figure 1b**).

For each pair of gene-wise parameters, we have

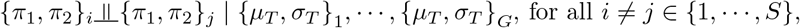

and similarly for each pair of sample-wise parameters, we have
 
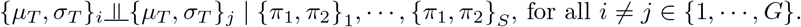

These relationships allow us to implement an optimization method, ICM, to iteratively derive the conditional modes of each pair of gene-wise or sample-wise parameters, conditional on the others [4]. Here, *π*_1_, *π*_2_ are constrained between 0 and 1, and *μ*_*T*_, *σ*_*T*_ are positively bounded. We combined a golden section search and successive parabolic interpolations to find a good local maximum [5] in each step. As shown by Besag [4], for ICM, the complete likelihood never decreases at any iteration and the convergence to the local maximum is guaranteed. Our ICM implementation is described in **Figure 5**.

**Figure 5.**
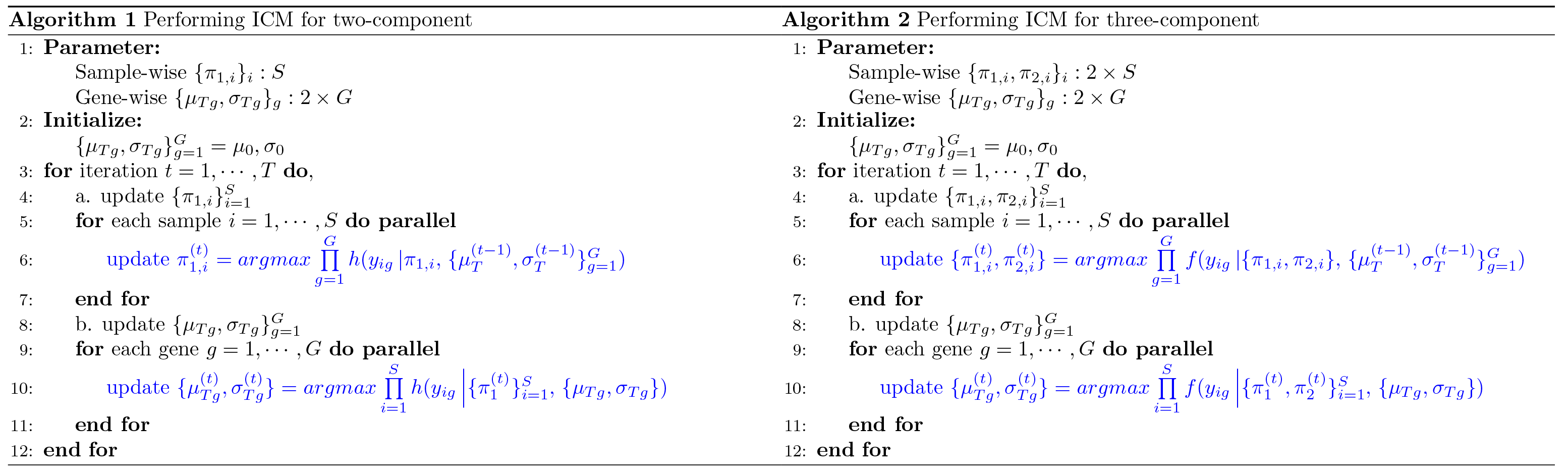
Outline of the ICM implementation in *DeMixT*. The *h*() represents the full likelihood based on a single integral for a two-component model; and *f*() represents the full likelihood based on a double integral for a three-component model.

### The GSCM approach to improve model identifiability

Due to the high dimension of the parameter search space, and often flat likelihood surfaces in certain regions of the true parameters (e.g., *μ*_1_ ≈ *μ*_2_) that will be encountered by ICM (**Supplementary Figure 2**), we have developed a GSCM approach (illustrated in **Figure 1b**) to focus on the hilly part of the likelihood space. This reduces the parameter search space and improves the accuracy and computational efficiency. Here, we describe our general strategy. As there are large variations in the number of genes that are differentially expressed across datasets, the actual cutoffs may be adjusted for a given dataset.

**Stage 1** We first combine the *N*_1_ and *N*_2_ components and assume a two-component mixture instead of three. This allows us to quickly estimate *π*_*T*_.

**a**: We select a gene set containing genes with small standard deviations (< 0.1 or 0.5) for both the *N*_1_ and *N*_2_ components. Among these genes, we further select genes with 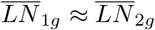 (mean difference < 0.25 or 0.5), where the 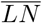 is the sample mean for the log2-transformed data. Within this set, we further select genes with the largest sample standard deviations of *Y*_*g*_ (top 250), suggesting differential expression between *T* and *N*.

**b**: We run *DeMixT* in the two-component setting to estimate *μ*_*Tg*_, 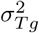 and *π*_*T*_.

**Stage 2** We then fix the values of {*π*_*T*_}_*i*_ as derived from Stage 1, and further estimate {*π*_1_}_*i*_ and {*π*_2_}_*i*_ in the three-component setting.

**a**: We select genes with the greatest difference in the mean expression levels between the *N*_1_ and *N*_2_ components as well as those with the largest sample standard deviations of *Y*_*g*_ (top 250).

**b**: We run *DeMixT* in the three-component setting over the selected genes to estimate *π*_1_ and *π*_2_ given *π*_*T*_.

**c**: We estimate the gene-wise parameters for all genes given the fixed *π*’s. Finally, given all parameters, per gene per sample expression level, *n*_1,*ig*_, *n*_2,*ig*_ and **t**_*ig*_ are reconstituted.

### Simulation study for the GSCM approach

To demonstrate the utility of GSCM for parameter estimation, we simulated a dataset with expression levels from 500 genes and 90 samples, 20 of pure *N*_1_-type, 20 of pure *N*_2_-type and 50 mixed samples. For the 50 mixed samples, we generated their proportions for all three components (*π*_1_, *π*_2_, *π*_*T*_) ~ *Dir*(1, 1, 1), where *Dir* is a Dirichlet distribution. For each mixed sample, we simulated expression levels of 500 genes for the *N*_1_ and T-component from a log_2_-normal distribution with *μ*_*N*_1*g*__ and *μ*_*T*_*g*__ from *N*_[0,+∞]_(7,1.5^2^), and with equal variance. For the *N*_2_-component, we generated *μ*_*N*_2*g*__ from *μ*_*N*_2*g*__ + *d*_*g*_, where *d*_*g*_ ~ *N*_[−0.1,0.1]_(0,1.5^2^) for 475 genes 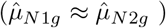 and *d*_*g*_ ~ *N*_[0.1,3]_(0,1.5^2^) ∪ *N*_[−3, −0.1]_(0,1.5^2^) for 25 genes 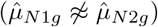. Then we mixed the *N*_1_, *N*_2_ and T-component expression levels linearly at the generated proportions according to our convolution model. We created a full matrix consisting of 20 *N*_1_-type reference samples (generated separately from the *N*_1_ distribution), 20 N_2_-type reference samples (generated separately from the *N*_2_ distribution) and 50 mixed samples at each simulation and repeated the simulation 100 times for each of the three variance values *σ* ∈ {0.1,0.3, 0.5} to finally obtain 300 simulation repeats. We first ran *DeMixT* with GSCM, where we used 475 genes with simulated 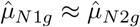 to run the two-component deconvolution (*N* versus *T*) and used the remaining 25 genes to run the three-component deconvolution with estimated 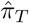. We also ran *DeMixT* without GSCM using all 500 genes. RMSEs between estimated and true proportions for all mixed samples were calculated for each of the two runs: with and without GSCM.

## Data analysis

All analyses were performed using the open-source environment R (http://cran.r-project.org). Documentation (knitr-html) of all scripts is provided at the GitHub repository.

### Summary statistics for performance evaluation

*Concordance correlation coefficient (CCC)*. To evaluate the performance of our method, we use the CCC and RMSE. The CCC *ρ*_*xy*_ is a measure of agreement between two variables, *x* and *y*, and is defined as 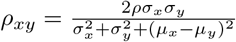, where *μ* and *σ^2^* are the corresponding mean and variance for each variable, and *ρ* is the correlation coefficient between the two variables. We calculate the CCC to compare the estimated and true proportions to evaluate the proportion estimation. We also calculate the CCC to compare the deconvolved and observed expression values (log_2_-transformed).

*Measure of reproducibility*. To assess the reproducibility of the estimated *π* across scenarios when the different components are unknown (i.e., three scenarios for a three-component model with one unknown component), we define a statistic 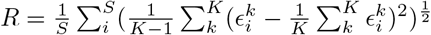, where 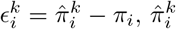 is the estimated value for the *k*-th scenario and *π*_*i*_ is the truth for sample *i*. *S* denotes the sample size and *K* is the number of scenarios. This measures the variations in the estimation errors across different scenarios. We consider a method with a smaller *R* as more reproducible and therefore more desirable.

### Mixed tissue microarray dataset

We downloaded dataset GSE19830 [21] from the GEO browser. We used the R package {*affy*} to summarize the raw probe intensities with quantile normalization but without background correction as recommended in previous studies [14]. We evaluated the performance of *DeMixT* with regard to tissue proportions and deconvolved expression levels on the set of genes that were selected based on the GSCM approach. Specifically, we selected genes with sample standard deviation < 0.1 in *N*_1_ and *N*_2_ components, among which we used those with 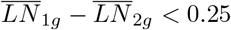 for running the 2-compoment model, and used the top 250 genes with largest 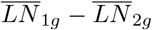 and largest sample standard deviation in *Y* for running the 3-component model. Then we ran ISOpure for the purpose of comparison.

### Mixed cell line RNA-seq dataset

This dataset was generated in house by mixing RNAs from three cell lines at fixed proportions. We mapped raw reads generated from paired-end Illumina sequencing to the human reference genome build 37.2 from NCBI through TopHat (default parameters and supplying the -G option with the GTF annotation file downloaded from the NCBI genome browser). The mapped reads obtained from the TopHat output were cleaned by SAMtools to remove improperly mapped and duplicated reads. We then used Picard tools to sort the cleaned SAM files according to their reference sequence names and create an index for the reads. The gene-level expression was quantified by applying the R packages GenomicFeatures and GenomicRanges. We generated a reference table from the human reference genome hg19 and then used the function findOverlaps to count the number of reads mapped to each exon for all the samples. This count dataset was pre-processed by total count normalization, and genes that contained zero counts were removed. The pre-processed count data were used as input for *DeMixT* and ISOpure.

We performed the same GSCM step as in the analysis of mixed tissue microarray data.

### Laser-capture microdissection (LCM) prostate cancer FFPE microarray dataset

This dataset was generated at the Dana Farber Cancer Institute (GSE97284 [24]). Radical prostatectomy specimens were annotated in detail by pathologists, and regions of interest were identified that corresponded to benign epithelium, prostatic intraepithelial neoplasia (abnormal tissue that is possibly precancerous), and tumor, each with its surrounding stroma. These regions were laser-capture microdissected using the ArcturusXT system (Life Technologies). Additional areas of admixed tumor and adjacent stromal tissue were taken. RNA was extracted by AllPrep (Qiagen) and quantified by RiboGreen assay (Life Technology). RNA labeling was performed using the SensationPlus FFPE method (Affymetrix) and hybridized to Affymetrix Gene Array STA 1.0.

FFPE samples are known to generate overall lower quality expression data than those from fresh frozen samples. We observed a small proportion of probesets that presented large differences in mean expression levels between the dissected tissues: tumor (*T*) and stroma (*N*) in this dataset (**Supplementary Table 7**). Only 53 probesets presented a mean difference 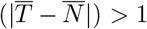, as compared to 10,397 probesets in GSE19830. We therefore chose the top 80 genes with the largest mean differences and ran both *DeMixT* and ISOpure under two settings: tumor unknown and stroma unknown.

### TCGA HNSCC data

We downloaded RNA-seq data for HNSCC from TCGA data portal (https://portal.gdc.cancer.gov/). There was a total of 44 normal and 269 tumors samples for HNSCC. We collected the information of HPV infection for the HNSCC samples. Samples were classified as HPV+ using an empiric definition of detection of > 1000 mapped RNA-seq reads, primarily aligning to viral genes E6 and E7, which resulted in 36 HPV+ samples [17]. We then devised a workflow to estimate the immune cell proportions (**Supplementary Figure 8**). Our workflow included three steps. The downloaded normal samples provided reference profiles for the stromal component in each step. We first downloaded stromal and immune scores from single-sample gene set enrichment analysis for all of our tumor samples [25] and selected 9 tumor samples with low immune scores (< −2) and high stromal scores (> 0), which suggested that these samples were likely low in immune infiltration. We then ran *DeMixT* under the two-component mode on these samples, generating the deconvolved expression profiles for the tumor and stromal components. We used these profiles as reference samples for running *DeMixT* under the three-component mode in the 36 *HPV*^+^ samples, generating deconvolved expression profiles for the immune component. In these two steps, we used deconvolved profiles that have smaller estimated standard variations as the reference profiles for the next step. We then ran *DeMixT* under the three-component mode on all 269 samples with reference profiles from normal samples and the deconvolved immune component. We calculated p-values (Benjamini-Hochberg corrected [3]) for the differential test of deconvolved expressions for the immune component versus the stromal component, and for the immune component versus the tumor component, respectively, on a set of 63 immune marker genes.

We performed gene selection in the GSCM approach (as described above), with a slightly larger threshold to account for the large sample size: sample standard deviation < 0.5 and the top 500 genes for three-component deconvolution to estimate the *π*’s.

## Ethics approval and consent to participate

Not applicable.

## Consent for publication

Not applicable.

## Availability of data and materials

1. The public data used in this study are GSE19830 [22] and GSE97284 [23] from GEO browser, and RNA-seqV2 count data from the Genomic Data Commons Data Portal [1].
2. The RNA-seq count data used for validation were generated from our lab and can be downloaded from https://github.com/wwylab/DeMixr. The FASTQ files will be uploaded to GEO browser upon publication.
3. The *DeMixT* source code and the entire analytic pipeline are available at https://github.com/wwylab/DeMixT.

## Author’s contributions

Z.W. developed and coded the algorithms in *DeMixT*, analyzed the data and performed the validation studies. S.C performed the application study and analyzed the data using *DeMixT*. J.A. proposed the assumptions of linearity and model distributions. F.G. and R.J. helped build the *DeMixT R* package. S.T., B.L., W.L., X.T., I.I.W., M.B, L.M. and M.L contributed data/materials for the validation and application studies. Z.W. and W.W. wrote the manuscript. J.S.M, S.T., G.P. and C.C.H. contributed to the discussion of results and revision of the manuscript. W.W. supervised the whole study. All authors read and approved the final manuscript.

## Acknowledgements

We thank Vesteinn Porsson, Ilya Shmulevich, David Gibbs, Liuqing Yang and Hongtu Zhu for useful discussions and valuable suggestions on the paper.

